# Genome size and repeat content contribute to a complex architecture of flowering time in *Amaranthus tuberculatus*

**DOI:** 10.1101/2023.07.13.548797

**Authors:** Julia M. Kreiner, Solomiya Hnatovska, John R. Stinchcombe, Stephen I. Wright

## Abstract

Genome size variation, largely driven by repeat content, is poorly understood within and among populations, limiting our understanding of its significance for adaptation. Here we characterize intraspecific variation in genome size and repeat content across 186 individuals of *Amaranthus tuberculatus*, a ubiquitous native weed that shows flowering time adaptation to climate across its range and in response to agriculture. K-mer based genome size estimates vary by up to 20% across individuals, with transposable elements, unknown repeats, and rDNAs being the primary contributors to this variability. The additive effect of this variation has important phenotypic consequences—individuals with more repeats, and thus larger genomes, show slower flowering times and growth rates. Compared to newly-characterized gene copy number and polygenic nucleotide changes underlying variation in flowering time, we show that genome size remains a modest but significant contributor to the genetic basis of flowering time. Differences in flowering time across sexes and habitats are not mirrored by genome size variation, but rather polygenic variation and a gene copy number variant within the ATP synthesis pathway. Repeat content nonetheless shows non-neutral distributions across the genome, and across latitudinal and environmental gradients, reflecting numerous governing processes that in turn influence quantitative genetic variation for phenotypes key to plant adaptation.

**Author Summary:** The remarkable and seemingly inconsequential variation in genome size across species has long been an enigma in evolutionary biology. Calling this viewpoint into question, correlations between genome size variation and traits linked to fitness are increasingly uncovered. While this suggests that DNA content itself may be a source of adaptive genetic variation, repeat elements that propagate at the cost of the host are known to largely mediate this variation and may thus limit adaptive potential. Here we look to disentangle these multi-level dynamics, characterizing repeat dynamics across the genome and among individuals across diverse collections of a widespread agricultural weed, linking repeat content to genome size variation, and characterizing the relative importance of its phenotypic consequences. In *Amaranthus tuberculatus*, we find non-neutral repeat distributions across individuals across the range, and while this repeat variation underlies both variation in genome size and flowering time, we show that it makes a relatively minor contribution to variation in a fitness-related trait across the landscape relative to monogenic and polygenic features. Together, this work broadens our perspective on the complex selective dynamics that govern intraspecific variation in genome size and traits key to fitness in plants.

## Introduction

The genome was traditionally viewed as a blueprint containing information to encode the phenotype; however, genome size and composition are also quantitative traits in themselves [1,2]. Early views on genome size evolution led to the prediction that more complex organisms, with a wider array of cell and tissue types, would have more complex genetic encoding—more genes and larger genomes. Yet, investigations of variation in genome size revealed a puzzling lack of correlation with perceptions of organismal complexity [3–5]; genome size was not in fact a clear predictor of ‘information content’. Variation in the amount of repetitive DNA sequences is now widely recognized as an important resolution to this paradox. In flowering plants, the primary driver of variation in repetitive content is transposable elements (TEs), which proliferate by creating copies of themselves in new sites across the genome, and whose proportional content is known to range between as much as 85% of the genome in maize and as little as 20% in *Arabidopsis thaliana* [6,7]. Nevertheless, the role of repetitive sequence in shaping genomic and phenotypic variation across populations remains unknown for most species.

A classic model explaining TE abundance variation involves a balance between the rate of transposition increasing TE insertions and the force of negative selection removing them due to their deleterious effects [8]. These effects stem from the ability of TEs to insert into and near genes, to cause ectopic recombination, and to affect transcription of nearby genes through the spread of epigenetic silencing [9–14]. On the one hand, if effective population size and thus the magnitude of drift and efficacy of selection differs among populations, population-level TE abundances may differ [15,16] and covary with population structure. Similarly, in genomic regions of low recombination (e.g., sex-determining regions), the efficacy of selection is reduced, driving another source of variation in TE content across the genome and between the sexes [17], which has recently been implicated in sex differences in survival [18,19]. On the other hand, TE abundance may differ among individuals and populations due to variation in transposition, as a consequence of TEs differentially evolving to evade host silencing mechanisms [20] or due to local physiological responses to environmental stress. Biotic and abiotic stressors, such as temperature, irradiance, nutrient starvation and fungal pathogens, are known to induce higher rates of transposition of certain TEs [21–24]. Finally, TEs may also be a source of local adaptation at multiple scales, by influencing the expression, copy number, and mutational landscape of particular focal genes (reviewed in [25,26]) and/or by generating variation in genome size that in turn influences quantitative traits [2].

While the evolutionary drivers of genome size variation are still debated [27], genome size is increasingly considered an important contributor to plant adaptation [28]. Larger genomes may not only mean a larger adaptive mutational target size (especially for non-genic regulatory variation; discussed in [28]) but may also have direct developmental consequences: larger genomes are often associated with larger cells, slower cell division, slower organismal growth rates, and ultimately a longer time to maturity and reproduction [29–32]. While the nature and direction of causality between genome size and these genomic properties has been difficult to uncouple [33], variation in genome size is hypothesized to be under selection as a source of variation for local adaptation via life history rate timing and is supported by recent observations of intraspecific variation in corn (*Zea mays)* [2,34]. At high altitudes, where faster growth and time to flowering assure reproduction before the early end of the season, maize individuals not only harbour fewer TEs, heterochromatic knobs, and smaller genomes compared to lower altitude plants but also show a faster rate of cell production and earlier flowering time [2]. The association between genome size and elevation described in [2] remained significant even after controlling for genome-wide relatedness, consistent with a model of selection acting on genome size through its effects on flowering time (or highly correlated traits). Taken together, variation in TE abundance across populations may be mediated not only by the balance between transposition rate and selection on individual elements, but also by spatially fluctuating selection on repeat abundance through its effect on life history traits.

To further evaluate how and why repeats vary in abundance within species, we turned to the prevalent agricultural weed, *Amaranthus tuberculatus* (common waterhemp). *A. tuberculatus* is an annual, dioecious, wind-pollinated plant, whose range is centered around the Mississippi river in the United States [35]. The species consists of two varieties, which were historically isolated to the northeast (var. *tuberculatus*) and southwest (var. *rudis*) of midwestern USA and southeastern Canada. Key to the dynamics of adaptation in the species is variability in life history traits (e.g., flowering time, growth rates) across geographic clines, habitats (natural and agricultural), and by varietal ancestry [36]. The species has evolved rapidly to industrial agriculture over the last six decades, driven by extreme selective pressures on genome-wide standing genetic variation drawn preferentially from southwestern var. *rudis* ancestry [37]. However, the contribution of variation in repeat content and genome size to these adaptive dynamics has yet to be explored.

Drawing on previously published high coverage (∼28x) genomic data from 187 individuals spanning natural and agricultural environments from Kansas to Ohio, accompanied by phenotypic measurements from a quantitative genetic common garden experiment, we address the following questions: (1) What is the distribution of repeat content across the genome and among individuals of *A. tuberculatus*? (2) Which repeat types are the predominant drivers of genome size variation and what are the consequences for key life history phenotypes? and (3) How important is genome size in encoding flowering time and responding to environment mediated selection? To do so, we combined k-mer-based genome size estimates, base pair abundances for 16 repeat and TE classes across individuals, and a novel characterization of the genetic architecture of flowering time. Our findings demonstrate the complexity of selective forces that govern variation in repeat abundance, genome size, and life history, and that interact to determine local adaptation and sex differences in this ubiquitous species.

## Results

### Transposable element variation across the genome

We first set out to characterize TE composition across the *A. tuberculatus* female reference genome [38]. TEs make up 66.28% or ∼443 Mb of the 668.5 Mb *A. tuberculatus* genome **(Fig 1A)**. In total we annotated 888,765 TEs, including two taxonomic repeat orders of cut-and-paste DNA transposons (366,445 terminal inverted repeat elements [TIRs] and 182,448 Helitrons) and two orders of copy-and-paste Retrotransposons (317,533 long terminal repeat elements [LTRs] and 1841 non-LTRs/LINEs). Mean lengths ranged from 316 and 329 bps for annotated Helitrons and TIR elements to 781 bps for LTR elements. The greatest proportion of the genome was represented by LTR elements (including unknown LTRs, Copias, and Ty3 elements) in part due to their larger size, as is typical in plants (reviewed in [39]). Helitrons in comparison were nearly twice in number as individual LTR element families but are at least half their size on average.

**Fig 1.**
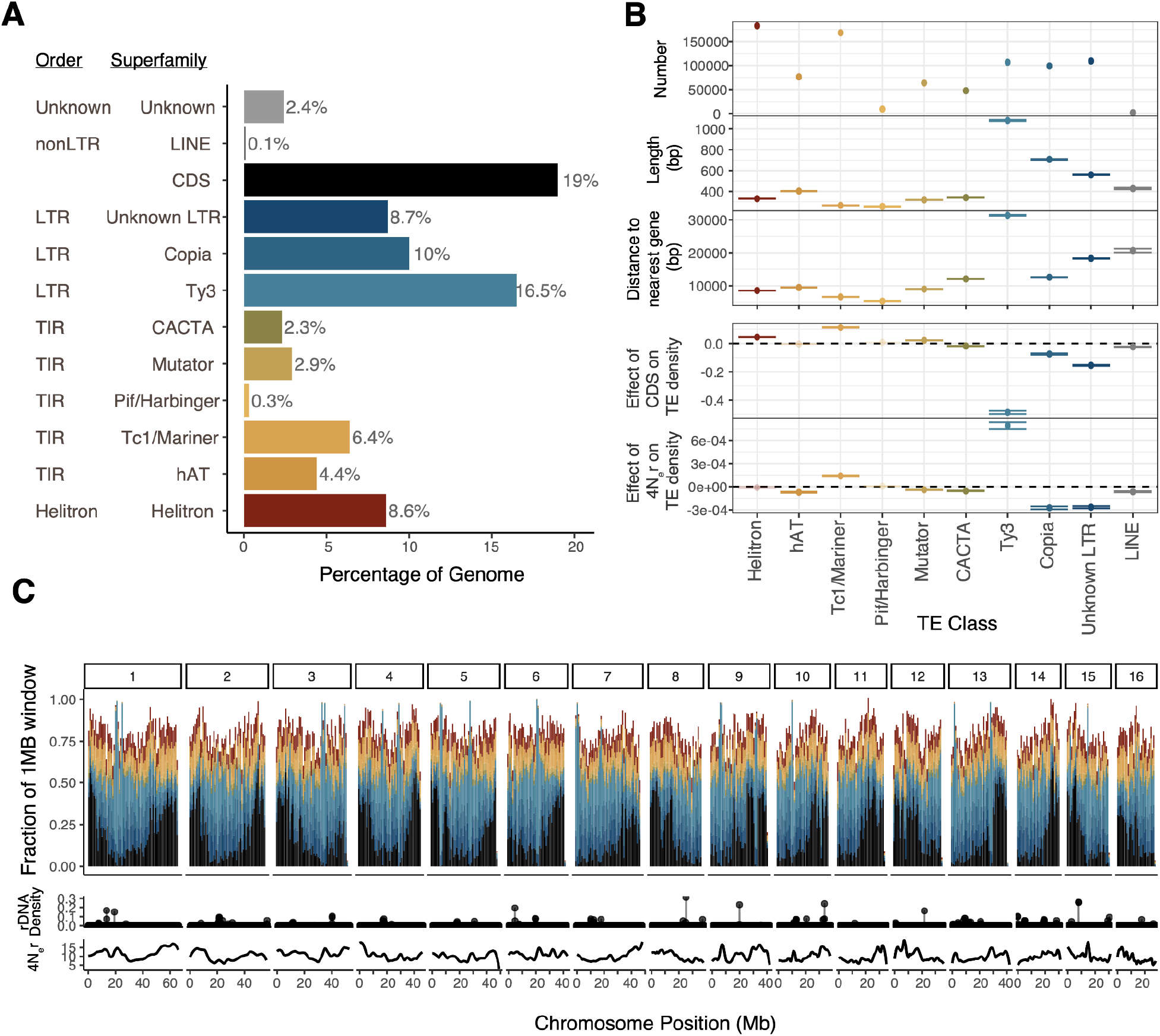
Variation in TE composition across the female *Amaranthus tuberculatus* reference genome by order and superfamily. **A)** The fraction of the female *A. tuberculatus* reference genome composed of different TE orders and superfamilies. **B)** Statistical summaries by TE superfamily, illustrating the differences in the number, size, and distance to genes across the reference genome. The bottom two rows represent the effect of coding sequence density (CDS) and *4N_e_r* on TE density as inferred from a multiple regression for each TE superfamily, for which all opaque lines are significant at p<0.05. Horizontal bars represent standard error of the estimate. **C)** The distribution of TE superfamilies and coding sequence content across the 16 chromosomes (top; colour codes from A and B), relative to the 100 kb window density of rDNAs (middle) and means of the 100 kb population scaled recombination rate (4N_e_r; bottom).

While TE superfamilies vary in size and number within *A. tuberculatus*, the distribution of TEs also varies across the genome in relation to genic content (coding sequence density; CDS) and the population recombination rate (4N_e_r) (the two of which are positively correlated in *A. tuberculatus*), likely reflecting differences in activity, transposition biases, and the strength of negative selection [17] (**Fig 1C, Sup Figs 1 & 2**). LTR elements were the largest and typically found furthest from genes, while TIRs (e.g., PiF/Harbingers, Tc1/Mariners) tended to be the smallest and closest to genes (Fig 1B). When we tested how well the population recombination rate and CDS density predicted the composition of TE superfamilies in 100kb windows in a multiple regression framework, we found every possible combination of the direction and strength of these predictors across TE superfamilies **(Fig 1B)**. Variability in LTRs across the genome was the most consistently explained by the strong negative effect of CDS (Copias: F=240, p < 10^-15^; Ty3s: F=1823, p<10^-15^, Unknown LTRs: F=1558, p <10^-15^), while typically exhibiting negative correlations with population recombination rate (Copias: F=238, p<10^-15^; Unknown LTRs: F=340, p<10^-15^; but negative for Ty3: F=356, p<10^-15^; **(Fig 1B)**. The correlations of population recombination rate and CDS were much more variable across TIR superfamilies, while variability in Helitrons is not explained by recombination rate but is positively correlated with CDS (F=211.8262, p < 10^-15^; **Fig 1B)**. Clearly, the *A. tuberculatus* genome reflects a complex genomic landscape of TE diversity as is seen in other systems (e.g. [40]).

**Fig 2.**
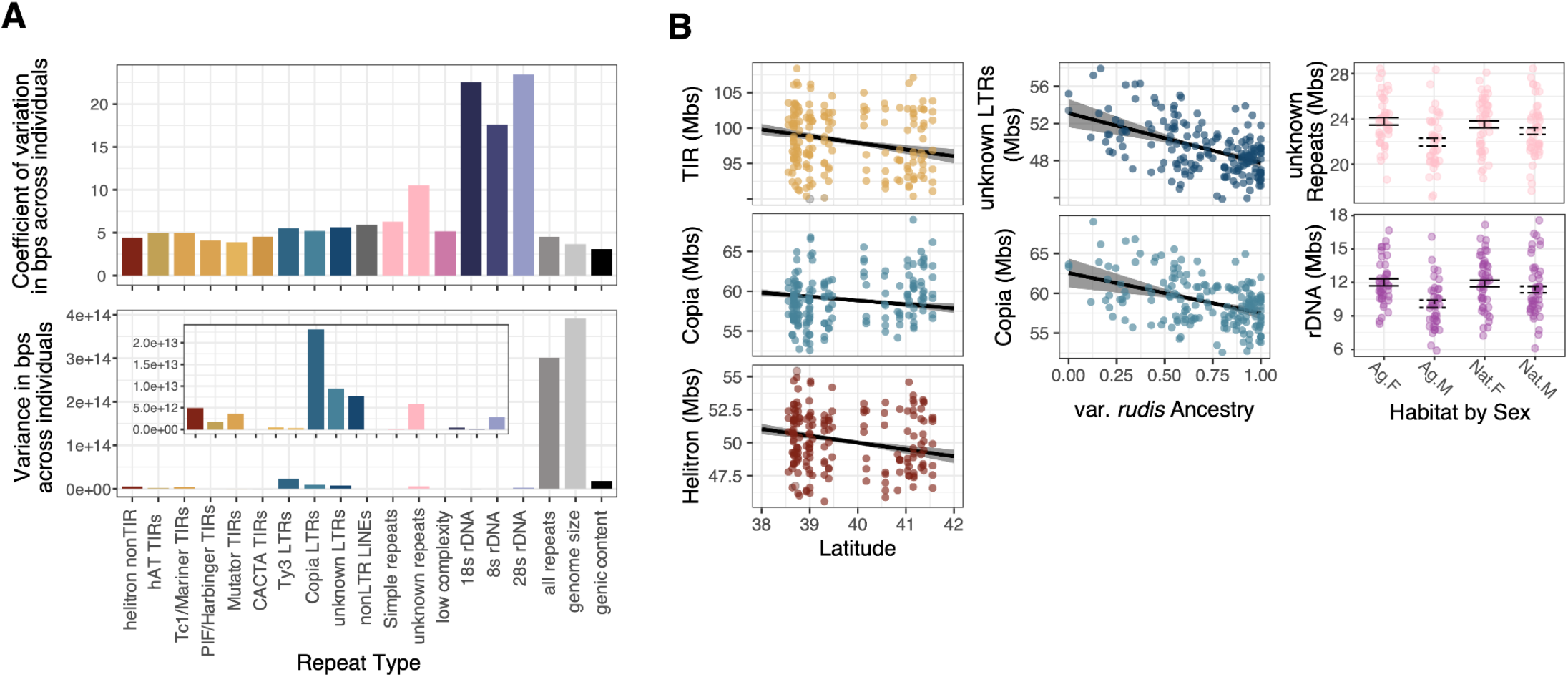
Variation in repeat abundance across individuals. **A)** The coefficient of variation (top) and variance (bottom) in bp amount of an individual’s genome composed of a given repeat class. **B)** The relationship between repeat classes with latitude (left), var. *rudis* ancestry (middle), and the interaction between habitat (Ag: agricultural site; Nat: natural site) and sex (right). Points represent raw data, while regression lines and error bars represent the least squares mean from a mixed effect model that accounts for relatedness. Only significant relationships shown.

We also annotated simple repeats and rDNA genes. rDNAs, repetitive genes functioning in ribosome production [41], made up 4.1% of the genome and were distributed across all 16 main chromosome scaffolds. A total of 2968 5S rDNA genes, 55 28S rDNA genes and 52 18S rDNA genes were annotated. Simple, low complexity, and unknown repeats comprise 7.0%, 1.5%, and 11.1% of the genome, respectively.

### Repeat variation across individuals

Looking beyond a single reference genome, we investigated the extent of intraspecific variation in TE and repeat class abundance in *A. tuberculatus* and thus the potential for it to contribute meaningfully to adaptive evolution. We estimated the abundance of each repeat class and TE superfamily (see methods; as delimited in Fig 1A) within the genomes of 187 individuals. The median bp composition of repeat classes across individuals showed approximately the same rank order as annotated in our reference genome: Ty3s (86.8 MB), Copias (59.0 MB), Helitrons (50.1 MB), and unknown LTRs (49.3 MB) show the greatest mean bp contribution across individuals, while 5s rDNA (1.24 Mb), low complexity repeats (0.86 Mb) and non-LTR LINEs show the least (0.58 Mb) **(Sup Fig 3 & 4)**. We next quantified both the variance and the coefficient of variation (CV, standard deviation scaled by the mean) in the bp composition of repeat types across individuals (**Figure 2A)**. The three rDNA subunits had the highest CV, more than double that of nearly all other repeat classes. By contrast, TEs tended to show the lowest coefficient of variation among individuals; there was proportionally little variability in abundance of TE classes (as measured by CV), although the absolute variance in abundance was high given the large number of these elements (particularly Ty3, Copia, and Unknown LTRs) (**Figure 2A).**

**Fig 3.**
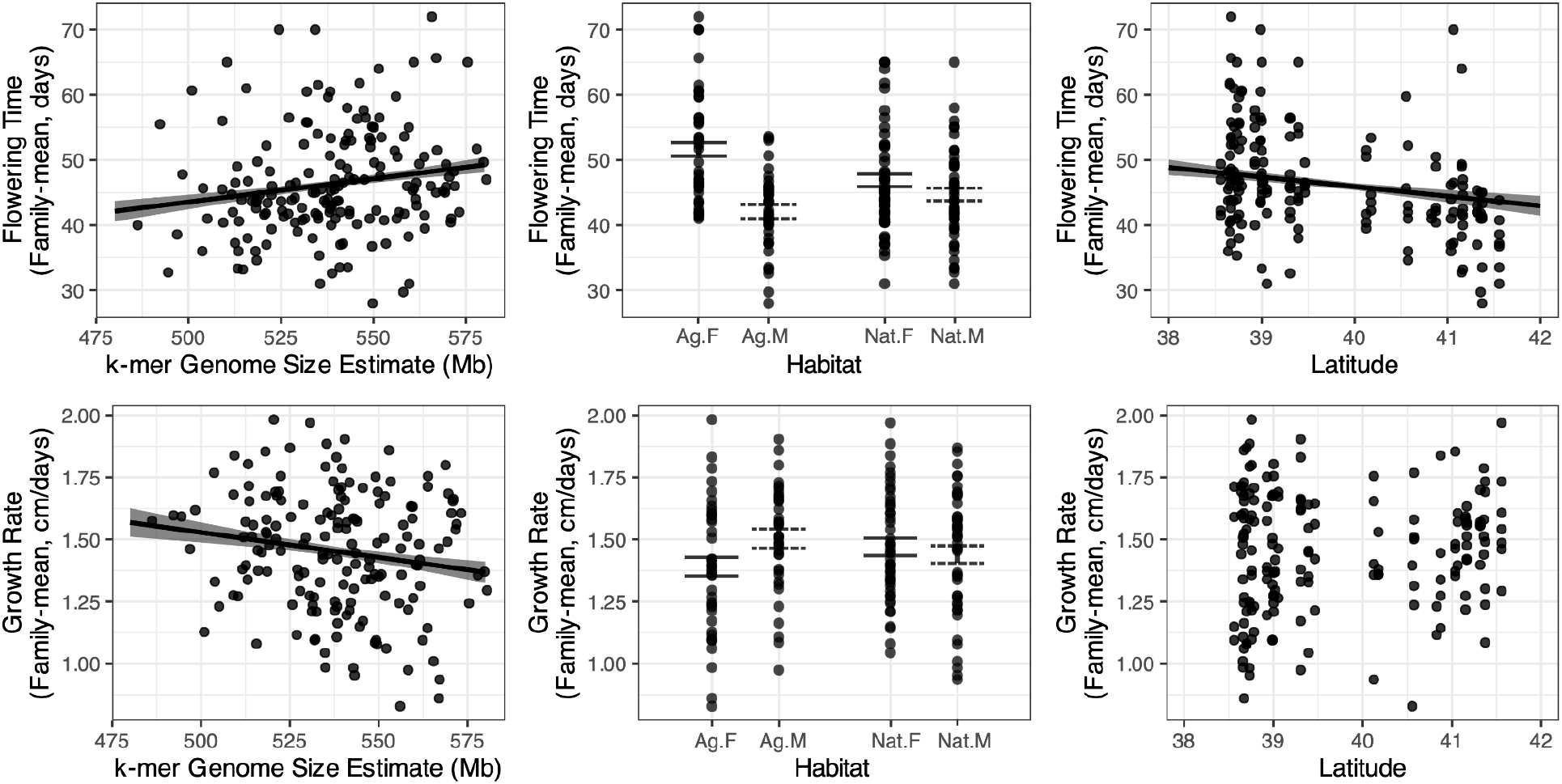
Genome size predicts flowering time (top) and growth rate (bottom) in *A. tuberculatus,* traits that also differ by habitat in a sex specific manner. Points show raw data, while regression lines and error bars depict least squares mean estimates from linear mixed modeling of flowering time and growth rate. Trend lines are shown for all significant relationships.

#### Landscape and organismal predictors of repeat content

We inferred landscape and organismal level predictors of each TE superfamily and repeat class abundance using linear mixed models that included the relatedness matrix (i.e., population structure) as a random effect (as in [2,42], with the matrix visualized in **Sup Fig 5**; see methods **eq. 4** for mixed model). In doing so, we found widespread evidence of latitudinal clines in repeat content that exceeded expectations from neutral population structure (visualized in **Sup Fig 1)**. The abundance of several elements declined substantially with latitude, including TIR elements (B=-942124, χ^2^ =5.41, p = 0.020) [Tc1: F= 7.44, p = 0.006; PiF: F=6.92, p= 0.009; hAT: F=4.57, 0.032; Mutator (F=5.22, p=0.022)], Helitrons (B=-516999, F=5.90, p=0.015), and Copia elements (B=-488909, F=3.89, p=0.049) **(Fig 2B)**. Because *A. tuberculatus* varietal ancestry varies along this same axis, ancestry proportion (based on structure inference, as in [36]) was explicitly included as a fixed effect predictor in addition to the relatedness matrix, suggesting that such latitudinal clines are not simply the result of the different timescales of evolutionary history. Ancestry did, however, significantly predict two repeat types, Copia elements (B=-5061330, F=3.94, p=0.047) and unknown LTRs (B=-5369193, F=6.41, p=0.011) (**Fig 2B)**, suggesting that a history of reproductive isolation and/or geographic variation in climate (or highly correlated [a]biotic forces) play a role in mediating these repeat abundances across the range.

We next investigated whether *A. tuberculatus*’ recent colonization of agricultural habitats was associated with the abundance of particular repeat classes. No repeat class showed evidence for an effect of current habitat (natural or agricultural). However, we did detect a significant sex by habitat interaction for two repeat classes (**Fig 2B)**, where differences in repeat abundance between sexes depended on whether the comparison is made within natural or agricultural habitats. For repeat classes that were significantly less abundant in males, rDNAs (sex effect: χ^2^=14.92, p=0.0001) and unknown repeats (sex effect: χ^2^=12.74, p=0.0004), the difference between sexes was greater in agricultural compared to natural habitats (sex by habitat effect; rDNAs: χ^2^=4.29, p=0.038; unknown repeats: χ^2^=3.28, p=0.07) **(Fig 2B)**.

### Variation in genome size

To understand the cumulative effects of this variation in repeat content, we next quantified genome size variation across individuals. Based on k-mer inference [43], there is substantial genome size variation across 186 individuals in this study, with the largest estimate being ∼20% larger than the smallest estimate (min Mb = 486.2, max Mb = 580.5, mean Mb = 538.8; **Sup Fig 6**, **Fig 3**), about double the variability seen in *A. thaliana* (up to 10% [1]). K-mer-based estimates are known to underestimate genome size and especially so for species with a past history of whole genome duplication [44,45], and indeed the average genome size estimate from these inferences falls outside of the confidence interval of genome size inferred from flow cytometry analyses of 3 individuals (95% CI 621.4–729.8 [46]). Nevertheless, our estimates of genome size correlate strongly with total repeat content in bps (r=0.83, p < 10^-15^) and, as expected, more weakly with gene content in bps (r=0.17, p = 0.024). The estimated coefficient of variation we observe for genome size is lower than any individual repeat type **(Fig 2A)**, likely reflecting the uncoupling of different repeat class abundances within and across individuals.

To test which repeat types contribute most to variation in genome size, we modeled genome size as a function of the different repeat types and genic content (methods eq. 1). When comparing the major categories of repeats including TEs, rDNA, simple repeats, low complexity regions and other unknown repeats as independent variables in such a model, TEs (F=195.85, p < 10^-15^), rDNAs (F=19.37, p=1.9 x 10^-15^), and unknown repeats (F=34.97, p=1.8 x 10^-8^) significantly explain genome size variation among individuals. The full model explains 72% of the variance in estimated genome size, with TEs alone explaining 32.3% of the variation in genome size, rDNAs 3.2%, and unknown repeats 5.8%.

### The phenotypic consequences of genome size variation

We leveraged phenotypes measured in a common garden experiment in these same samples [36] to test whether genome size correlates with key life history traits. Focusing on traits related to growth and the timing of key life history transitions, we modeled the effects of genome size on growth rate (the increase in plant height between the 4-6 leaf stage and flowering) and time to flowering. Because sequenced individuals spanned multiple treatments in the common garden experiment [36], here we used family-mean phenotypes as measured in the control treatment. With individuals collected from a broad sample across habitats (natural and agricultural), latitudes, longitudes, varietal ancestries, and with separate sexes in the species, we included all such factors along with genome size as fixed variables in this model of flowering time, and the relatedness matrix as a random effect (**methods eq. 2)**. Furthermore, with differences in sexual dimorphism across environments having been identified in other wind pollinated species [47], we tested for sex by habitat interactions.

We found that genome size is a significant predictor of flowering time (F =7.70, p=0.006), with every additional 10 MB predicted to delay flowering by 0.7 days (Figure 3C), in a mixed effect model. In our collections, that corresponded to approximately a week difference in flowering time between samples with the smallest and largest sampled genome. Genome size explained 4.4% of the variation in flowering time in this model. Overall, sex explained the most variation in this model of flowering time (partial r^2^ = 20%, F=41.83, p =1.05×10^-11^) with males flowering 9.55 days [SE = 1.47 days] earlier than females. The second strongest predictor of flowering time was a sex by habitat interaction (F = 13.52, p=0.0002) reflecting greater sexual dimorphism in flowering time in agricultural compared to natural habitats (**Fig 3B**). We also found a main effect of habitat (F=11.29, p=0.0007), longitude (F = 7.69, p=0.0056), and latitude (F = 5.482, p=0.0192) (Fig 3A; as described in [36]). In a model of growth rate, genome size is positively related to growth rate and the most significant predictor (F=4.57, p=0.034; **Fig 3C**), explaining 2.8% of the variation in growth rate, similar to the effect of sex (partial r^2^ =2.7%, F= 4.42, p=0.037). We also find a marginal habitat by sex interaction effect (F=3.85, p = 0.052) (**Fig 3B**). Taken together, these results support the hypothesis of genome size playing a role in determining key life history traits in *A. tuberculatus*.

While flowering time, growth rates, and individual TE classes show significant latitudinal and environmental variation, we do not find any significant correlation between genome size and environment (latitude: ꭓ2 = 2.6267, p = 0.1051; habitat: ꭓ2 = 0.0012, p= 0.9728) or sex (ꭓ2 =0.0004, p = 0.9843), when accounting for neutral population structure through a relatedness matrix (methods eq. 3). Total genome size has apparently not responded directly to selection through flowering time, but instead varies as a product of underlying repeat dynamics.

### The relative importance of quantitative genome size variation, oligogenic, and polygenic features to flowering time evolution

We next characterized the contribution of large-effect and polygenic variants to these fitness related traits, ranging from copy number variation to SNPs. While genome-wide associations for SNPs and genic copy number variation with growth rate yielded no significant SNPs after Bonferroni or any level of FDR correction, flowering time showed multiple types of genetic associations.

A GWA of flowering time using gene-level copy number variation as predictors identifies 1,680/30,771 genes with a significant effect at the FDR q<0.05 level and 34/30771 genes at the Bonferroni p<0.05 level. These 34 genes include FLOWERING LOCUS D on Scaffold 11, with the broader set of genes being significantly enriched for the ATP synthesis PANTHER pathway (Bonferroni p-value=4.6 x 10^-3^), GO molecular functions in NADH dehydrogenase (Bonferroni p-value = 6.1 x 10^-6^) and proton-transporting ATP synthase activity (Bonferroni p-value = 6.9 x 10^-4^), as well as numerous other related GO biological functions (**Sup Table 1)**. This signal of enrichment appears to be predominantly driven by one large-effect locus on Scaffold 10 **(Fig 4A)**, a region in which a cluster of 7 NADH-ubiquinone oxidoreductases (two ND1, ND4L, ND5, ND6, two ND2), one NADH dehydrogenase (NAD7), 4 ATP synthases (ATP6, ATP MI25, atpA, atpB), and one Cytochrome b6-f complex subunit 5 (petG) map. A regression of flowering time on the gene with the strongest statistical association in this region (and genome-wide) reveals that 20% of the variation in flowering time can be explained by copy number variation at this locus, which varies from ∼2 to 14 **(Figure 4A)**.

**Figure 4.**
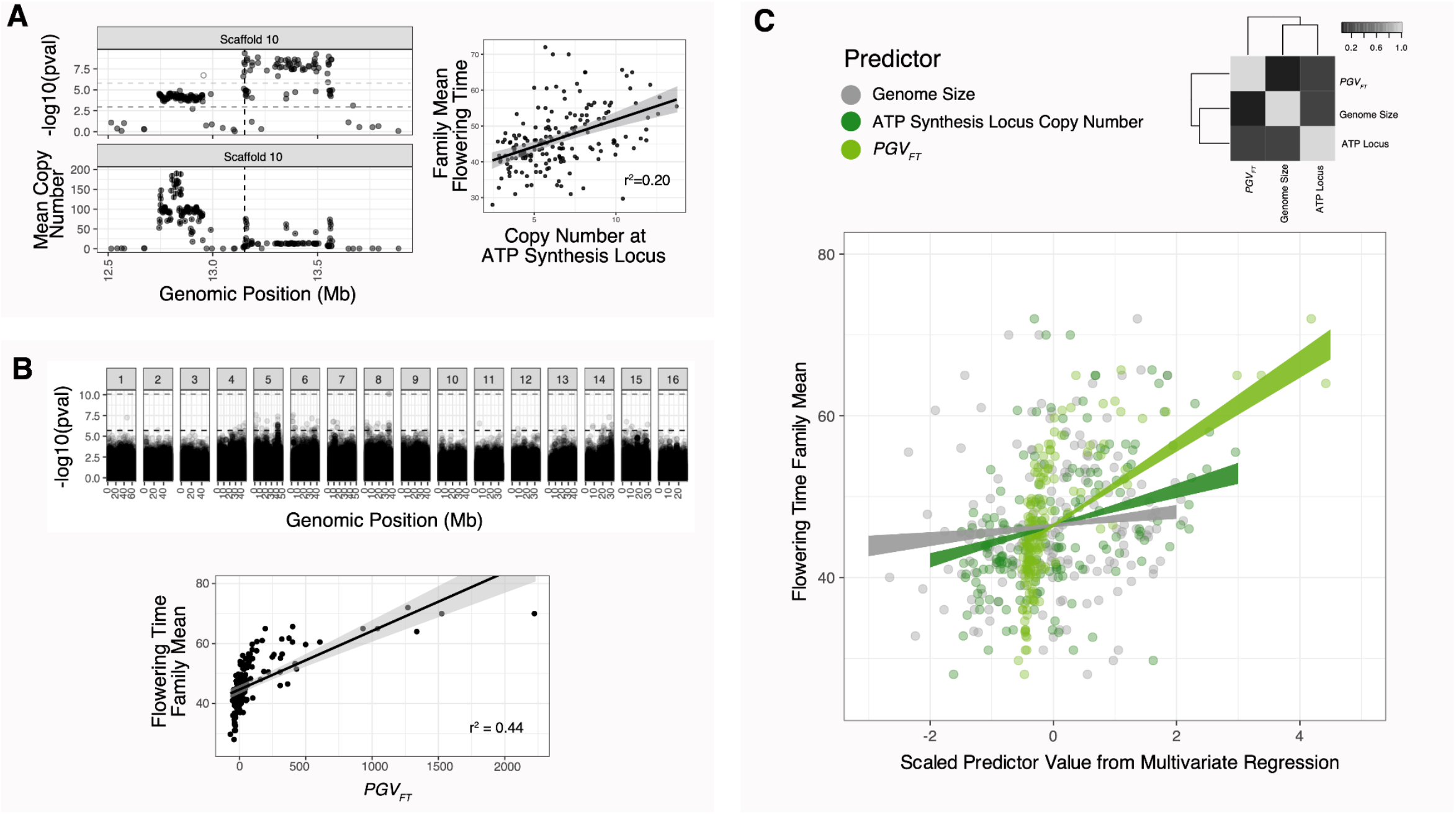
The genetic architecture of flowering time in *A. tuberculatus* and the relative importance of associated genetic features. **A)** The association of a copy number variant in the ATP synthesis pathway with flowering time (vertical line denoting locus with the most significant association genome wide). **B)** The polygenic value of individual flowering time (*PGV_FT_*) based on 96 SNPs that pass a 10% FDR correction from a genome-wide association (lower black horizontal dashed line, Bonferroni threshold also shown above). Black line in the bottom plot represents the linear regression fit between flowering time and *PGV_FT_*. **C)** A mixed effect model for flowering time while controlling for relatedness demonstrates the relative importance of associated genomic features, from the polygenic value in B) and copy number variation at the ATP synthesis locus in A) to genome size variation. Correlation structure of fixed effect predictors also illustrated in the top right of C).

**Table 1.** Biological GO Enrichment results after Bonferroni correction from the gene-level copy number GWAS.

| GO biological process complete | # | expected | Fold Enrichment | bonferroni p-value |
| --- | --- | --- | --- | --- |
| electron transport chain | 15 | 3.51 | 4.27 | 0.0219 |
| generation of precursor metabolites and energy | 38 | 11.45 | 3.32 | 0.00000234 |
| cellular metabolic process | 371 | 270.23 | 1.37 | 1.55E-09 |
| metabolic process | 489 | 377.73 | 1.29 | 2.73E-11 |
| cellular process | 556 | 447.08 | 1.24 | 3.26E-11 |
| cellular respiration | 17 | 4.57 | 3.72 | 0.0291 |
| energy derivation by oxidation of organic compounds | 18 | 5.01 | 3.6 | 0.0249 |
| photosynthesis | 26 | 7.26 | 3.58 | 0.000286 |
| translation | 38 | 15.52 | 2.45 | 0.00383 |
| macromolecule metabolic process | 327 | 249.74 | 1.31 | 0.0000472 |
| organic substance metabolic process | 447 | 353.81 | 1.26 | 2.82E-07 |
| organic substance biosynthetic process | 182 | 131.88 | 1.38 | 0.0213 |
| biosynthetic process | 193 | 142.48 | 1.35 | 0.0321 |
| peptide biosynthetic process | 39 | 15.69 | 2.49 | 0.00203 |
| peptide metabolic process | 41 | 16.63 | 2.47 | 0.00124 |
| organonitrogen compound metabolic process | 243 | 186.24 | 1.3 | 0.0171 |
| nitrogen compound metabolic process | 342 | 265.11 | 1.29 | 0.0000788 |
| amide metabolic process | 47 | 20.81 | 2.26 | 0.002 |
| amide biosynthetic process | 42 | 17.71 | 2.37 | 0.0026 |
| cellular nitrogen compound biosynthetic process | 100 | 55.01 | 1.82 | 0.0000637 |
| cellular nitrogen compound metabolic process | 210 | 137.15 | 1.53 | 7.31E-07 |
| cellular biosynthetic process | 148 | 97.37 | 1.52 | 0.00138 |
| organonitrogen compound biosynthetic process | 90 | 52.52 | 1.71 | 0.00385 |
| gene expression | 88 | 43.97 | 2 | 0.00000652 |
| primary metabolic process | 378 | 292.63 | 1.29 | 0.00000377 |
| nucleobase-containing compound biosynthetic process | 41 | 19.09 | 2.15 | 0.0374 |
| nucleobase-containing compound metabolic process | 159 | 107.14 | 1.48 | 0.00174 |
| heterocycle metabolic process | 176 | 126.82 | 1.39 | 0.0241 |

In contrast, SNP-level associations with flowering time appear to reflect a more dispersed, polygenic architecture. A GWA of flowering time using SNPs as predictors while controlling for the relatedness matrix identifies 96 loci in 73 genes across the genome passing a 10% FDR correction. We therefore calculated the polygenic value [48,49] for flowering time 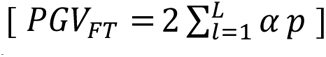, where *L* equals the 96 loci with an FDR q<0.1, *α* the effect size of a locus on flowering time, and *p* an individual’s allele frequency at that locus (genotype). A bivariate correlation of individual *PGV*_FT_with flowering time shows that the *PGV*_FT_explains 44% of the variation in flowering time in the environmental conditions in which it was measured (**Figure 4B)**.

The positive correlation of genome size and flowering time as previously described could also reflect the greater opportunity for particular phenotype affecting insertion events rather than a direct effect of repeat abundance on flowering time. To explore this hypothesis, we conducted a GWA of flowering time for repeat (as opposed to gene) copy number. The associations across the genome greatly mirrored that of the gene level analysis (Sup Fig 7) with only one clear peak on Scaffold 10 at the same ATP synthesis locus. While TEs in the region may have causally impacted flowering time, another likely explanation is that TEs were amplified along with numerous genes in this region, hitchhiking along with the phenotypic effects of the host genes. By jointly modeling the effect of copy number variation at this locus along with genome size on flowering time, we further distinguish these alternatives.

Finally, we tested the relative importance and independence of these polygenic and oligogenic predictors compared to genome size in a mixed effects model of flowering time that includes the relatedness matrix as a random effect. This model explains 95% of the variation in family-mean flowering time, 24% of which is attributed to the fixed-effect terms and 71% of which can be attributed to the genome-wide relatedness. Genome size remains a significant explanatory variable (ꭓ^2^ =4.10, p = 0.043) explaining 2.3% of the variation in flowering time but is considerably less important compared to copy number variation at the ATP synthesis locus (partial r^2^ = 12.8%, ꭓ^2^ =25.62, p = 4.2 x 10^-7^), which in turn is less important than *PGV*_FT_(partial r^2^ = 44.9%, ꭓ^2^ =162.6, p < 10^-15^) (**Fig 4C)**.

In contrast to genome size, these polygenic and oligogenic architectures show patterns of spatial differentiation reflecting selection on flowering time. Copy number variation at the ATP synthesis locus demonstrates significant variability among sexes (ꭓ^2^= 22.59, p = 2.01 x 10^-6^), across natural and agricultural habitats (ꭓ^2^= 9.27, p = 0.0023), and among sexes depending on habitat type (ꭓ^2^= 8.83, p = 0.0029), exceeding neutral expectations (methods eq. 3). Similarly, *PGV*_FT_shows a strong sex effect (ꭓ^2^=9.13, p =0.0025), a habitat effect (ꭓ^2^=4.16, p = 0.041), and marginally, a longitude effect (ꭓ^2^=3.06, p = 0.08).

## Discussion

We report marked intraspecific variability in repeat content and genome size that is associated with flowering time variation in *A. tuberculatus*. Individuals with more repeats and larger genomes tend to show slower growth rates and time to flowering. We leveraged past whole genome sequencing and common garden phenotype data for nearly 200 individuals to show that this quantitative variation in genome size complements polygenic and copy number variation for flowering time and is independent from the effects of locus-specific TE copy number on flowering time. In comparison to newly identified polygenic variation and a photosynthesis-related large effect gene copy number variation, genome size remains a significant but modest predictor of common-garden-measured flowering time across our collections.

Phenotypic and latitudinal associations with repeat content and/or genome size have been found across several systems, from maize [2,34,50] to *Drosophila* [51]. Mechanistically, cells with larger genomes are thought to take longer to undergo cell division and thus development, and this is supported by associations with cell size [32,52], cell production rate [2], stomatal density [32], flowering time [2], development time [53], growth form [54], and scaling laws [55]. We therefore predicted that earlier flowering waterhemp plants at higher latitudes, and in natural, not agricultural, habitats, might have smaller genomes and less repeat content. While we find evidence for an important effect of genome size on life history traits–both flowering time and growth rate–genome size does not vary across latitude and habitat beyond neutral expectations from genome-wide relatedness. This suggests that genome size predominantly impacts variation for flowering time within populations in *A. tuberculatus*, although this may also simply result from the minor contribution of genome size to flowering time. In part, the lack of signal of flowering time selection shaping genome size variation may result from the lack of large effect genome-size alleles in *A. tuberculatus*, such as heterochromatic knobs in maize [2]. Nonetheless, particular repeat classes do show clinal variation and differences across sexes, highlighting the numerous processes that are likely affecting the abundances of individual element families in *A. tuberculatus*.

Previous work in *A. tuberculatus* has revealed the sundry of genetic mechanisms underlying agricultural adaptation, from standing genetic variation and de novo origins to and gene flow [36,37,56]. Here we show the genetic architecture of flowering time appears to nearly as multifaceted. In addition to a role for genome size, we describe two genetic features underlying genetic variation for flowering time. We find copy number variation at a cluster of genes in the ATP synthesis pathway that is associated with variation in flowering time, adding to the increasingly recognized importance of structural variation in flowering time evolution [57–59]. That copy number variation for genes in the photosynthetic pathway predicts flowering time supports the notion that modifying photosynthesis can impact developmental rate and plant yield (e.g., [60–62]), providing some of the first evidence for such a link in natural populations. We also quantify polygenic variation for flowering time encoded by single nucleotide polymorphisms at 96 associated loci across the genome. These SNPs mapped to genes with molecular and biological functions varying from stigma and gynodiecium development (*HEC1*), meristem and flower development (*PCN, DOT2;* [63]), gibberlic acid mediated anther and seed development (*SPL8;* [64]), to DNA methylation and post transcriptional gene silencing (*AGO4, ROS1;* the former protein family having been implicated in mediating flowering time by modifying the expression of *FT* [65]) (**Sup Table 2**). Unlike genome size, both gene copy number and polygenic variation for flowering time showed differentiation among habitats and sexes, suggesting that strong selection for high performing, earlier flowering males [66–69] in agricultural habitats has been key to shaping the distribution of this genetic variation across the landscape.

**Table 2.**
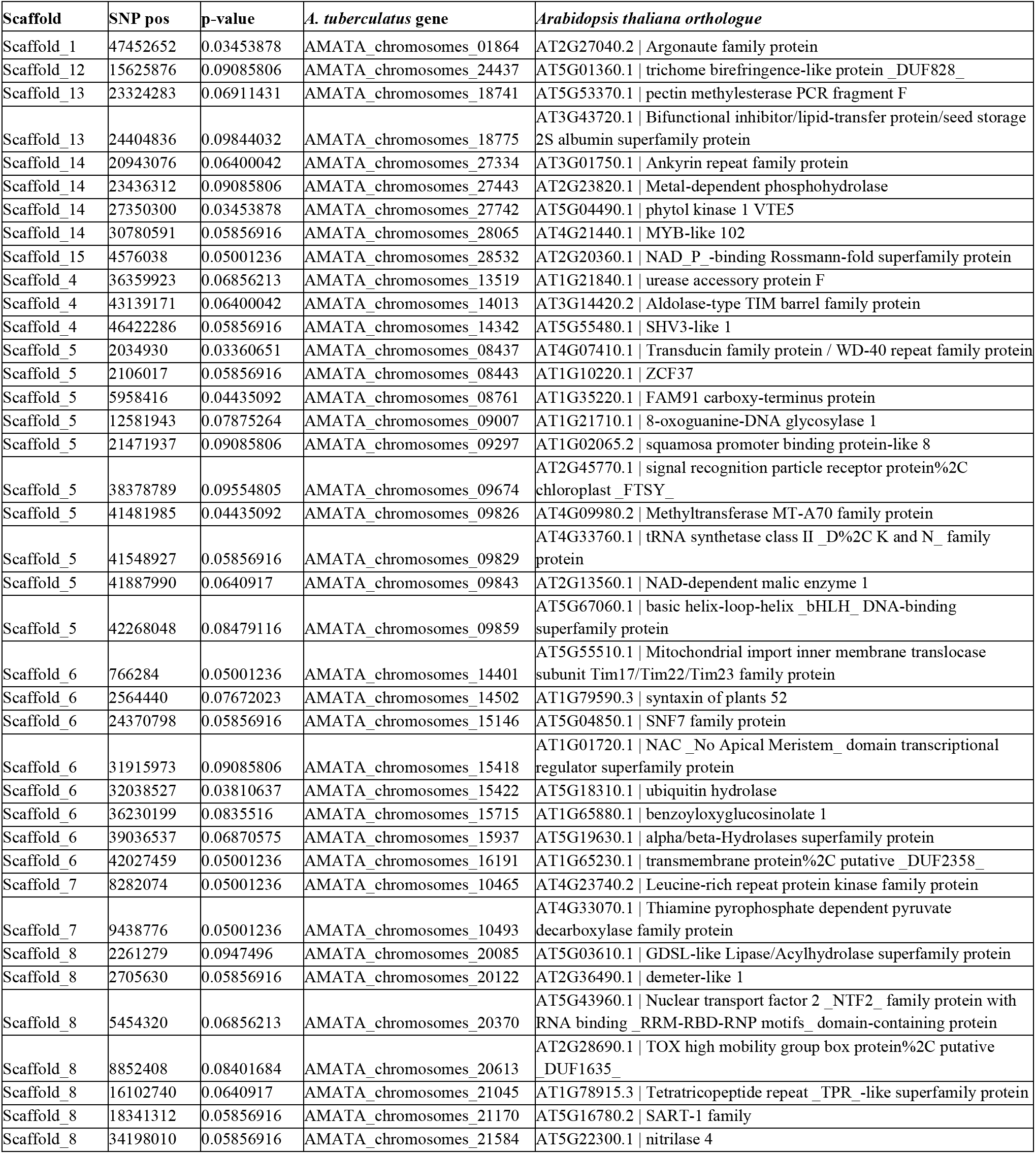
Significant SNPs from a GWA with flowering time, for those where their *A. tuberculatus* containing gene has an orthologous match in *Arabidopsis thaliana*.

While genome size does not show substantial variation among populations, the composition of repeat content does, suggesting the possibility of stabilizing selection on genome size with repeats competing for limited space in the genome [70–72]. The processes governing non-neutral latitudinal clines in TIRs, Helitrons, and Copia beyond neutral expectations are unclear, but could be mediated by differences in transposition rate and/or selection at the host or repeat level [73]) and may depend on their distribution across the genome. TE families varied tremendously in their correlations with gene density and the population-scaled recombination rate (4N_e_r). Recombination rate has been shown to vary with temperature [74–76] and thus latitude, implying that geographic co-evolution of repeat content and recombination rate [17] may occur to differential extents across repeat types based on their distribution across the genome. In *A. tuberculatus*, LTR density is strongly governed by CDS density, consistent with their enrichment in gene-poor centromeric regions across species [77] and a stronger selection for removal and/or an insertion preference away from genes in low recombination rate regions [17]. In contrast, TIR superfamilies are more often positively associated with CDS and more typically occurring in high recombination rate regions, which may drive their differential associations with latitude. Furthermore, methylation has been shown to covary with latitude and climate variables [78], with demethylation having a potentially adaptive role at low temperatures [79,80], suggesting that individuals with a higher content of active TE families may in part be due to differences in host silencing across environments. Overall, these results demonstrate that repeat associations with latitude and habitat are mediated by factors other than simply through their contributions to genome size.

TEs not only showed broadscale clines across the range, but across habitats (natural versus agricultural) depending on sex. We expect that linked selection played a key role in this finding. Recent work has shown that the *A. tuberculatus* chromosome that contains the male sex determining region harbours a fragment of the *Flowering Time* and *Heading date 3a* [81] in addition to the evidence we provide here of copy number and polygenic variation for flowering time being differentiated across sexes. Because the sex-determining region represents a large region of low recombination, one possibility is that agricultural selection on a haplotype that contained an early flowering variant of these genes could have by chance driven lower repeat content to higher frequency in such environments.

Taken together, the processes generating and governing variation for growth related life history traits are more complex than typically assumed. Flowering time is influenced by genome size—which, in the absence of polyploidy events and large-scale heterochromatic knobs, may predominantly reflect the balance between transposition and host removal for individual TE families, with some role for stabilizing selection on total genome size. We show this balance varies across numerous genomic, organismal, and environmental axes, and will thus require diverse range-wide collections (e.g. [82,83]) along with experimental quantification of transposition rates (e.g. [84]) and fitness effects to fully disentangle. Furthermore, quantification of individual element frequencies using long-read population sequencing will enable the characterization of element insertion frequencies, to assess the potential role of individual TE insertions in rapid adaptation to agricultural environments.

## Methods

We used the female reference genome as described in Kreiner et al., (2019). Briefly, the *A. tuberculatus* reference genome was sequenced and assembled from an individual female plant from an agricultural habitat. The resulting 2514 contigs were scaffolded onto a chromosome resolved reference genome of a closely related species, *A. hypochondriacus* [85], creating a reference with 16 pseudochromosomes [56]. The accompanying gene annotation (described in [56]) was also used.

For all analyses, we used the 182 samples which were previously sequenced and analyzed in [36,37]. Briefly, each sample comes from intra-population crosses that were performed to control for maternal phenotypic effects, with no two samples having the same maternal or paternal genotype. The collections originated from 17 paired agricultural and natural populations, all of which were located within 25 km of each other, and spanned three degrees of latitude and 12 degrees of longitude [36]. Out of 200 samples from unique maternal lines, 187 were successfully sequenced with short-read Illumina sequencing depths ranging in coverage between 20-35 X [36]. Due to a high error rate (>5%) in five of the samples were excluded from the study, leaving 182 samples for the estimation and analysis of repeat content. Phenotypic data for 176 of the sequenced samples, that among other traits, included flowering time and vertical growth rate was collected in a common garden experiment [36]. For each maternal line, 30 replicates were grown, one sibling in each of the three treatments (water supplemented, control, and soy competition), replicated across 10 experimental blocks in the common garden experiment [36]. For this study, we only used phenotypic data collected in the control treatment [36]. For each maternal lineage, flowering times were averaged across the ten siblings in the different experimental blocks. The proportions of var. *rudis* ancestry were estimated with the Faststructure algorithm using SNP genetic information, also described in [36,86]. The relatedness matrix from genome-wide SNPs across all individuals was computed in gemma [87] using the centered genotype matrix algorithm (**Sup Fig 5)**. Genome size was estimated by first counting 21-mers using the program KMC [88], and then by fitting a Bayesian model to the histogram of 21-mers using Genomescope2.0 [43].

### Non-overlapping TE annotation

EDTA (Extensive de-novo TE Annotator)-v1.9.7 was used to detect TEs in the reference genome and produce both a TE library and a non-overlapping annotation [89]. For TE library curation, the EDTA pipeline combines a number of high-performing TE finding programs and filters their outputs to produce a comprehensive and non-redundant TE library. LTRs are identified and filtered based on structural features by a combination of LTR_HARVEST_parallel, LTR_FINDER_parallel and LTR_retreiver [89–91]. Helitrons were detected by HelitronScanner-v1.1 which uses a two-layered local combinational variable (LCV) algorithm [92]. TIRs are detected by machine learning with the TIR-Learner2.5 program [93]. Following a number of filtering steps, the EDTA program reduces interlibrary redundancy between LTR-RTs, Helitrons and TIR elements, combines then into one library and clusters the TEs into families based on a modified 80-80-95 Wicker rule, resulting in one representative sequence per family. The resulting library of representative TEs is used to mask the genome with RepeatMasker-4.1.1 and the remaining unmasked regions are searched by RepeatModeler-2.0.1 for missed TEs, including SINEs and LINEs [94,95]. Redundancy between the RepeatModeler library and the EDTA library is then removed and the two libraries are merged to create the final EDTA TE library. Furthermore, any repeats that are detected but not identified as TEs by the EDTA program, such as simple repeats, are listed as repeat regions, which I refer to as unknown repeats in this study.

We then used the EDTA annotation function to annotate the *A. tuberculatus* reference genome. The EDTA annotation function combines the high confidence structure-based annotations produced by structure-based programs in EDTA with homology-based annotations produced by RepeatMasker-4.1.1, using the EDTA library. Additionally, the annotation resolved overlapping regions using the following priorities: structure-based annotation > homology-based annotation, longer TE>shorter TE> nested inner TE> nested outer TE [89].

### Ribosomal RNA annotation

RNAmmer -1.2 was used to annotate eukaryotic rDNA [96]. RNAmmer annotates rDNA using hmm’s that have captured structural features of rDNA from multiple alignments of rDNA database sequences across different species. The output included subunits: 28S, 18S, and 8S. The 8S subunit is most likely synonymous to the 5S subunit, as confirmed by doing a blast search of the identified 8S subunit, finding it aligns with >95% identity to 5S subunits in other plant species. For this reason, we refer to 8S rDNA as 5S rDNA in this study.

### Simple sequence annotation

To annotate simple sequences, low complexity regions and tandem repeats, we ran RepeatMasker on the reference genome with default parameters [94].

### Estimating repeat abundances

To estimate the abundances of TEs, rDNA, simple repeats and low complexity regions, we first created a nonoverlapping annotation file combining the TE annotation, rDNA annotation and the simple repeats annotation. The following order of priorities was used to resolve the overlapping sequences of repeats: rDNA>known TE>simple repeats and low complexity regions>unknown repeats. Any sequences that were left with fewer than or equal to 20 base pairs in length were removed, as they are more likely to be false positives. The non-overlapping annotation was then used to estimate both the copy number of individual TEs annotated in the reference for each individual, and the additive bp contribution of repeat classes to each individual’s genome. To do so, we calculated the mean coverage of reads mapping within each repeat using mosdepth [97] and scaled it by the median genic coverage genome wide to get an estimate of diploid copy number at each annotated region. We then multiplied the repeat element copy number by the length of that repeat, and summed within superfamilies and orders to get the bp repeat abundance for each, within each individual.

### Statistical analyses of genome size and TE abundances

The ggplot2 package was used for visualization in most figures [98], in combination with the plot_grid function in cowplot [99] to create panels of multiple figures. The size and number of TEs was visualized directly from the non-overlapping annotation based on counts and bp ranges of each superfamily. Distance to the nearest gene was calculated using bedtools closest [100], finding the gene closest to each repeat element, reporting the distance with option -d. The fraction of each genomic window composed of different repeat types was calculated with the bedops command ‘bedmap’ [101], using the commands --echo --count --bases-uniq-f. Population recombination rate was estimated with LDhat [102] in [56].

The variance and coefficient of variation of repeat abundance were both calculated in R, the latter by dividing the standard deviation by the mean of the repeat abundance distribution and multiplying by 100. To test which repeat types contribute most to variation in genome size, we implemented a model using “lme4” version 1.1-27.1 with genome size as the dependent variable and all the major categories of repeat types as the independent variables. Since genome sizes were estimated using a K-mer analysis, we weighed the samples by their fit in the K-mer estimated model:

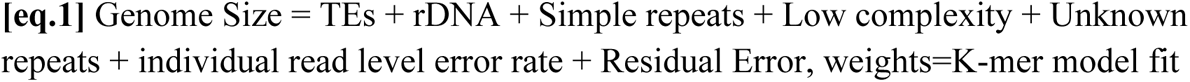

Percent of model variance explained by each term was calculated by comparing the r^2^ of the full model to a model reduced by each term. Note that predictors in this model were kept broad (i.e. TEs instead of Helitrons + LTRs + TIRs etc.) as a result of strong collinearity between repeat classes within individuals.

To test whether flowering time and growth rate can be explained by variation in genome size and which repeat abundances are driving this variation, we used the package lme4qtl that implemented a mixed effect model with the relatedness matrix as a random effect [103]. We also tested the relative importance of genome size on flowering time by including it in a model with two other genetic predictors of flowering time (copy number of an ATP synthesis pathway locus and the polygenic value for flowering time; **Eq 3**):

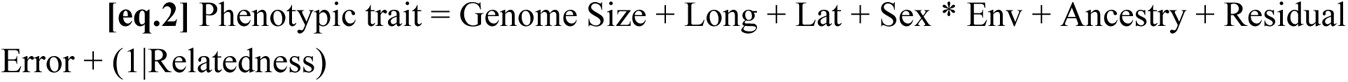

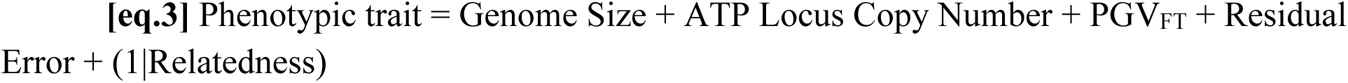

where the (1|) notation indicates a random effect after the bar operator. Percent of model variance explained by each term in the model was calculated with r2beta [104] and r2 for random versus fixed effects with the function r.squaredGLMM [105].

Similarly, we explored the predictors of individual repeat class abundances across populations. For each of the 16 repeat and TE classes (e.g. Copia LTRs, Ty3 LTRs, helitrons, 5S rDNAs, simple repeats, etc.), we implemented a mixed effect model with lme4qtl that controls for the relatedness matrix (as above), and that included the fixed effect terms:

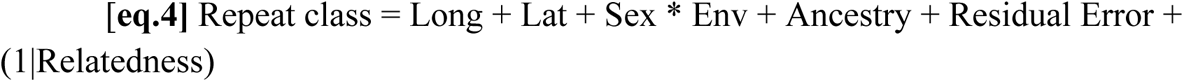

All models were evaluated with an anova using type III sums of squares.

### Characterization of other architectures underlying flowering time

We implemented two genome-wide association (GWA) approaches to better understanding the genetic architecture of flowering time in *A. tuberculatus*. Firstly, we investigated the extent that copy number variation is associated with flowering time through a GWA, using the family mean flowering time (and growth rate, to no avail) measured in the control environment from the common garden experiment in [36]. To estimate copy number variation, we used mosdepth [97] to get the mean read coverage within each gene for each individual, and then scaled it by an individual’s median genic coverage genome-wide to get an estimate of gene copy number. We used this gene by individual level information as the input for a GWA of gene copy number variation with flowering time. We applied the same coverage-based approach from the gene copy number GWA to implement a TE and repeat copy number variation GWA with flowering time. In both cases, we implemented a p-value FDR correction in R using the base function “p-adjust” and further investigated genes and repeats passing the 10% FDR threshold.

We next ran a GWA on high quality filtered SNPs from [36]. We did so using gemma [87] after using plink to convert from a vcf to binary file format, and using the gemma generated relatedness matrix as a covariate. With the 97 SNPs found across the genome that pass the 10% FDR threshold, we calculated each individual’s polygenic value [48,49] for flowering time. The polygenic value was calculated as:

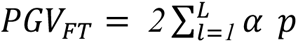

where *L* equals the 97 loci with an FDR q<0.1, *α* the effect size of the minor allele on flowering time from the GWA, and *p* an individual’s genotype (equal to 0.5 for heterozygotes and 1 for homozygous alternates). Our calculation of polygenic values for flowering time was based on genotypes at just the 97 SNPs genome-wide that passed a 10% FDR correction rather than genome-wide SNPs, in an effort to circumvent issues with uncontrolled population structure [106–108].

## Acknowledgements

Thanks to the Whitlock lab, Sally Otto, and Tyler Kent for feedback on the manuscript. JK was funded by a Killam Postdoctoral Fellowship and a Biodiversity Research Centre Bioinformatics Postdoc, S.I.W. was supported by an NSERC Discovery Grant and a Canada research chair. J.R.S. was supported by an NSERC Discovery Grant.

## Supplementary Figures and Tables

**Sup Fig 1.**
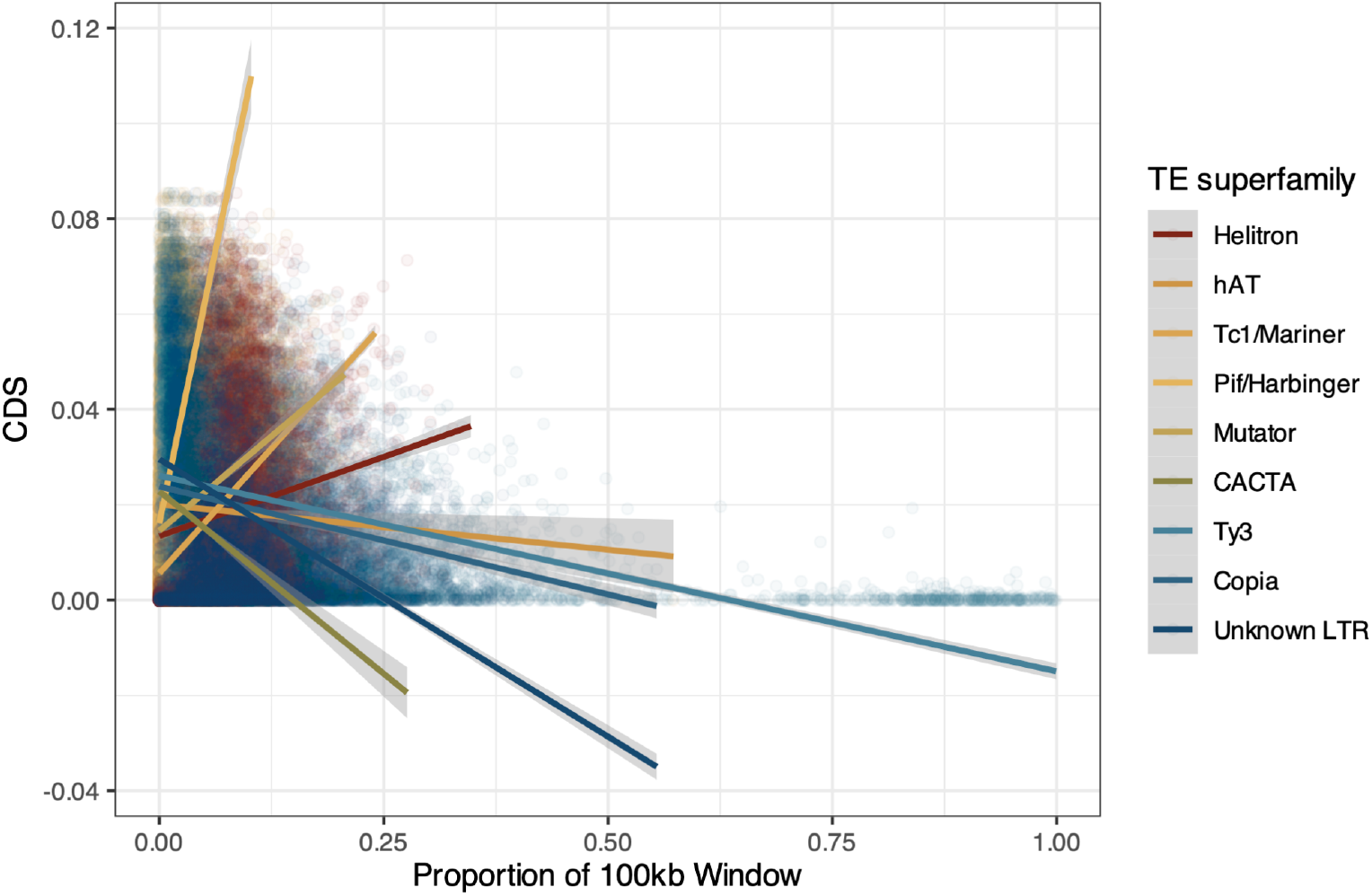
Bivariate correlations of TE density across the genome and coding sequence density across the genome.

**Sup Fig 2.**
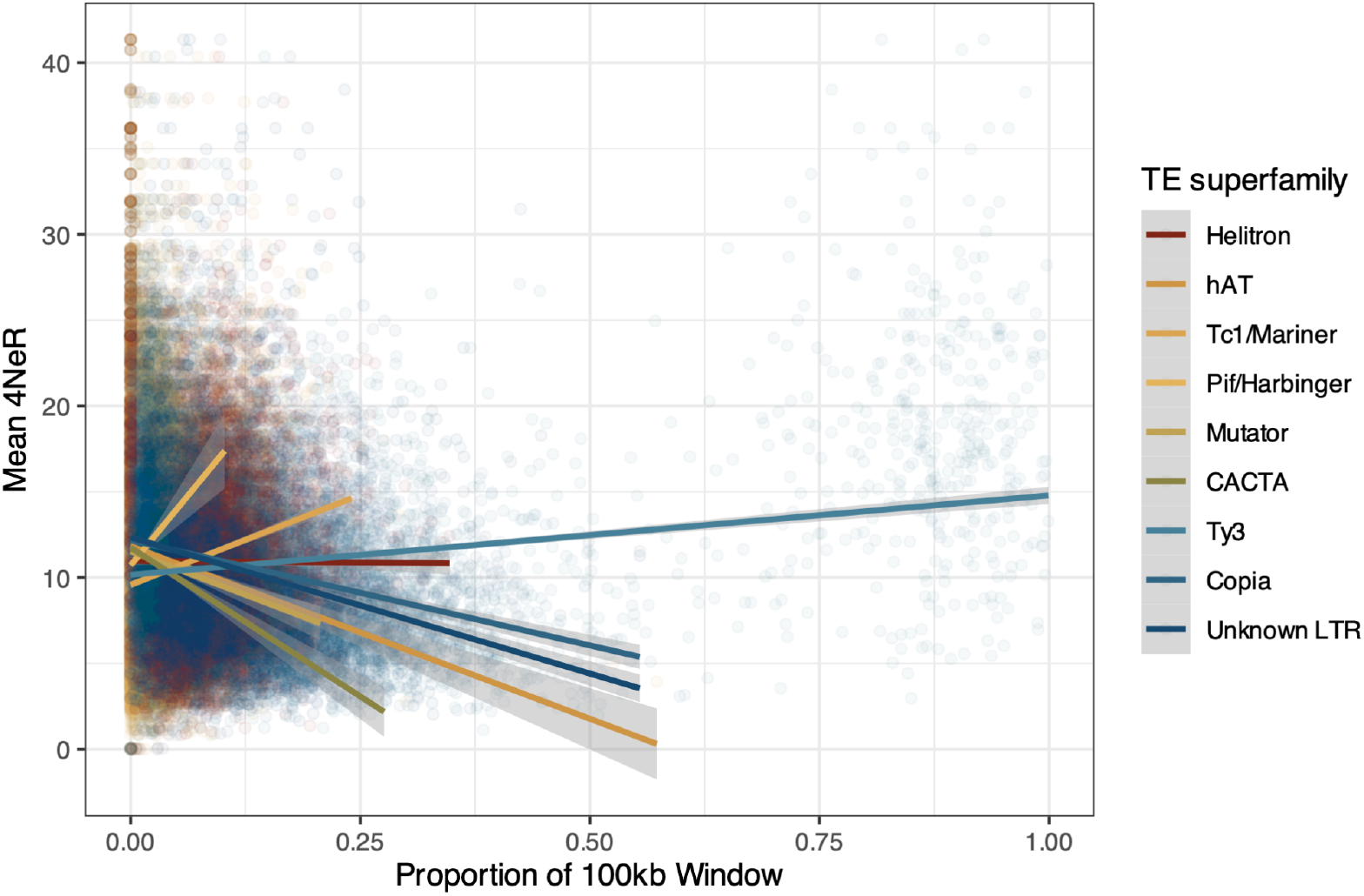
Bivariate correlations of TE density across the genome and coding sequence density across the genome.

**Sup Fig 3.**
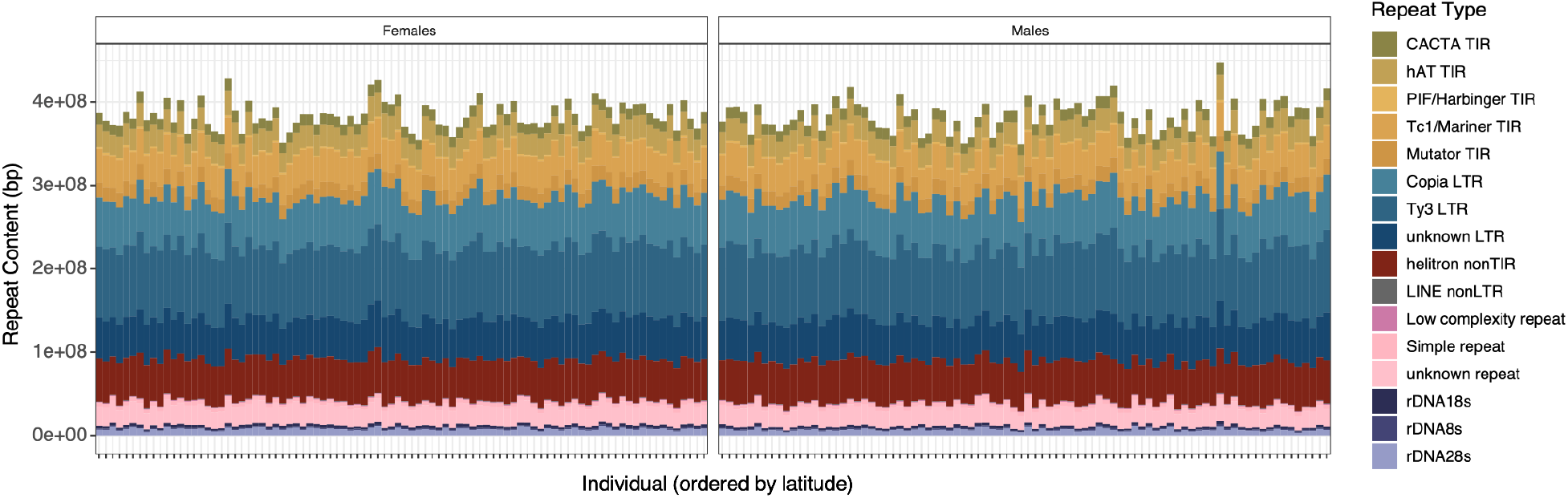
Repeat composition within and across individuals, with individuals oriented from south to north from left to right.

**Sup Fig 4.**
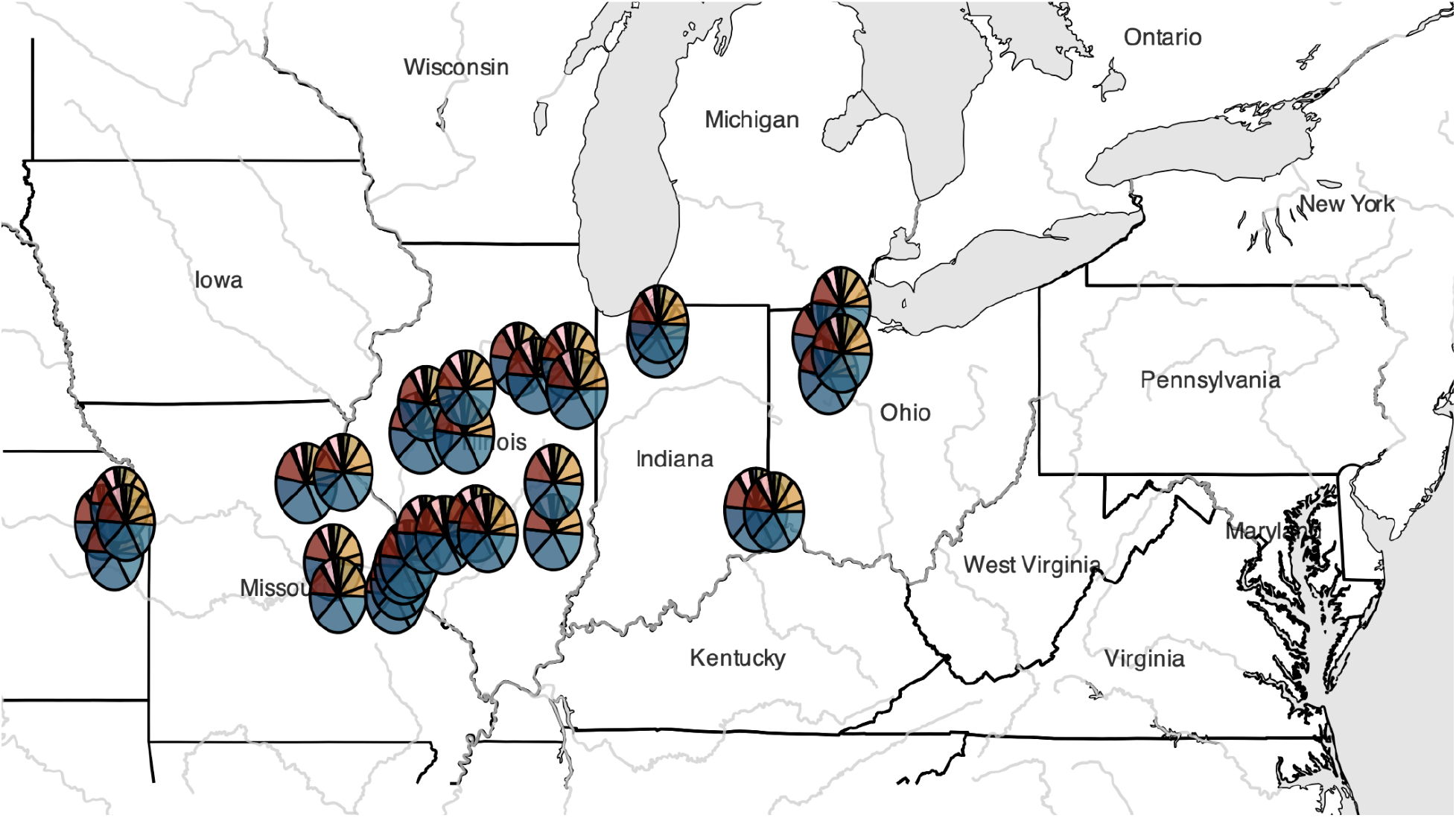
Mean repeat composition across populations of *A. tuberculatus*

**Sup Fig 5.**
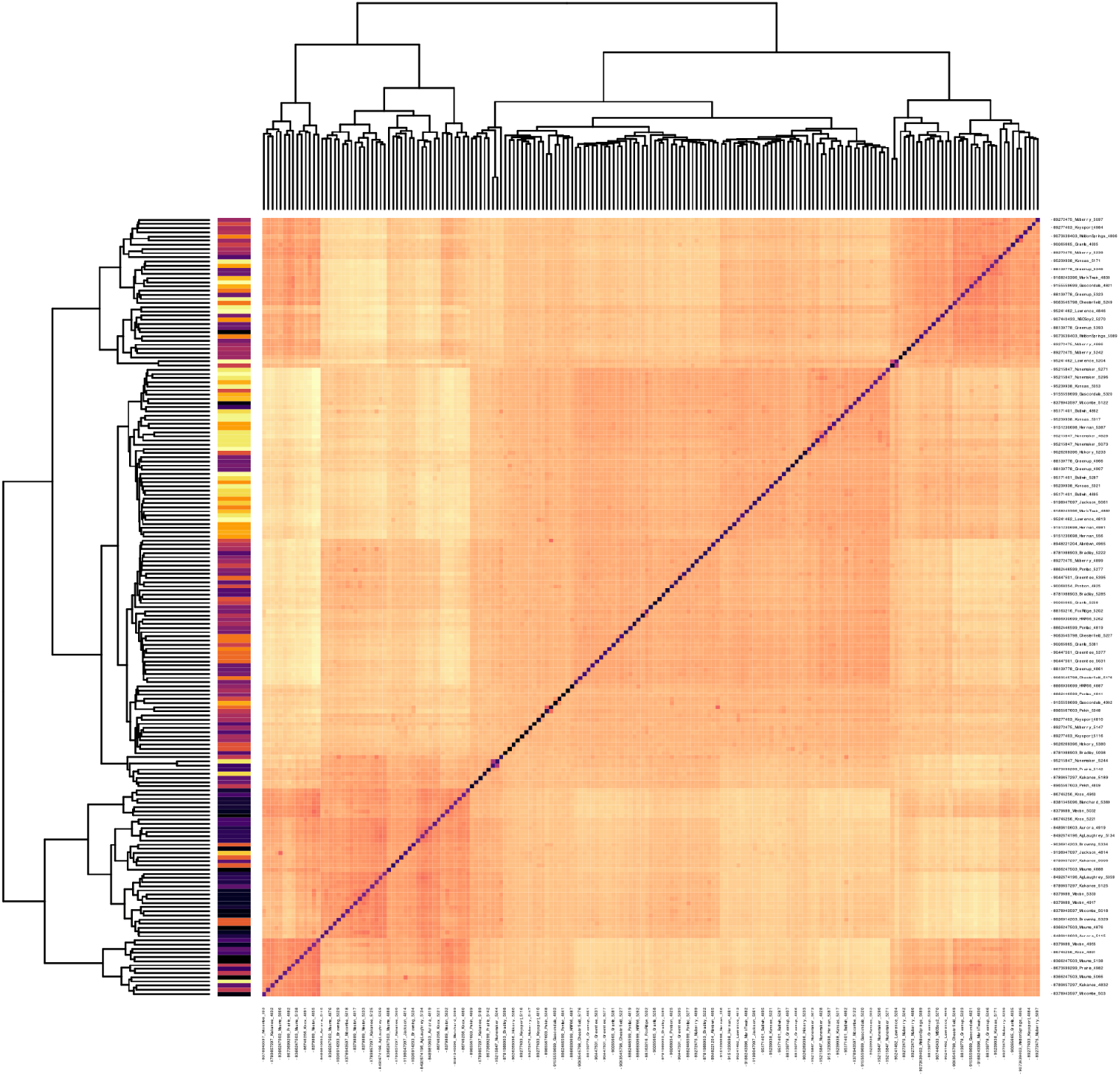
The relatedness matrix based on centered genotypes as computed in plink. Colors on the left represent population groupings, ordered by longitude (with the most eastern populations in darker colours).

**Sup Fig 6.**
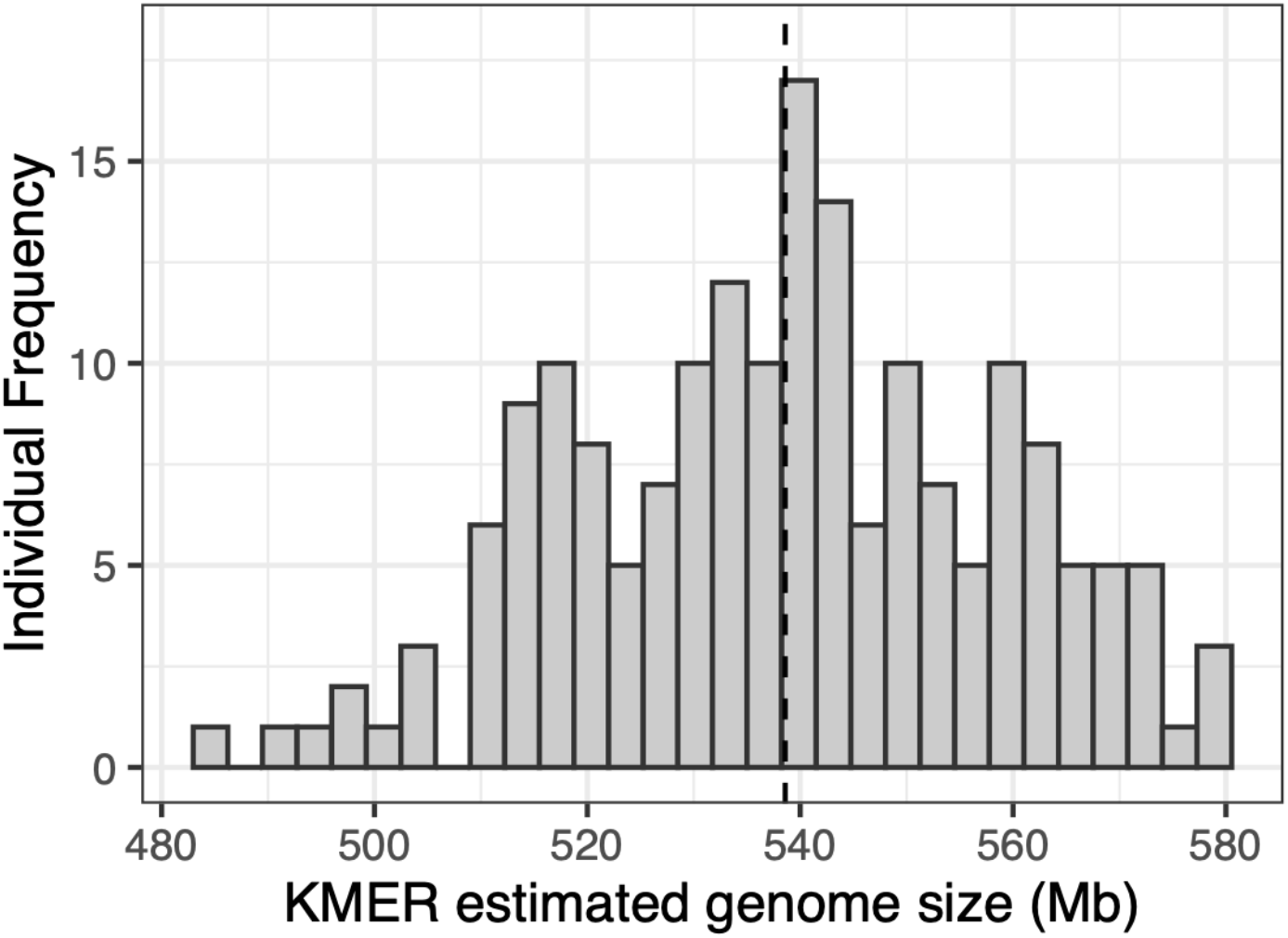
The distribution of k-mer inferred genome sizes in this study, where the mean estimated genome size is indicated by the vertical dashed line.

**Sup Fig 7.**
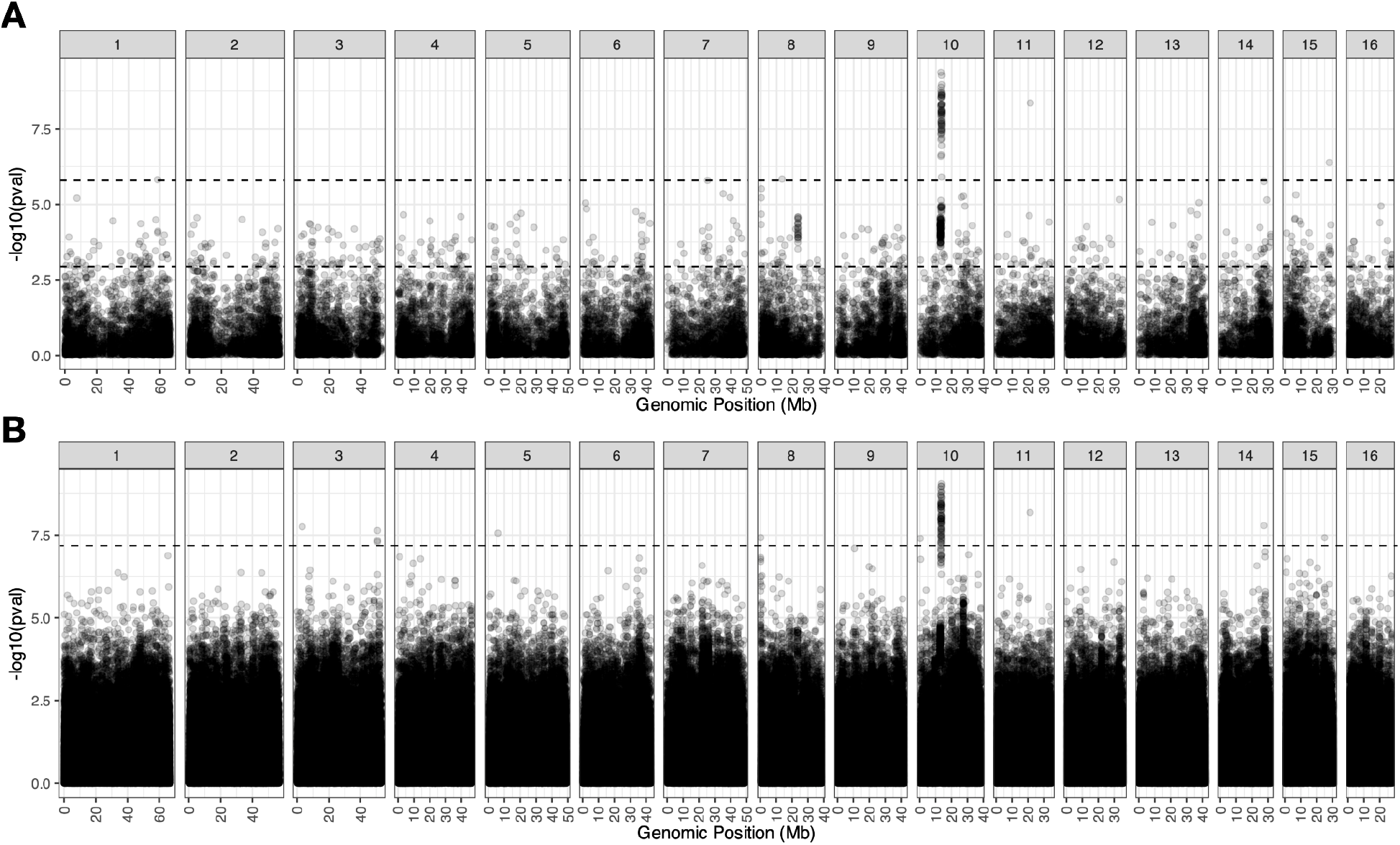
Copy number associations with flowering time across the genome, for **A)** genes, and **B)** repeats. Horizontal dashed line indicates a 5% false discovery rate threshold.

